# Hybrid conferences: opportunities, challenges and ways forward

**DOI:** 10.1101/2022.03.18.484941

**Authors:** Eleonora Puccinelli, Daniela Zeppilli, Paris Stefanoudis, Annaïg Wittische-Helou, Marjorie Kermorgant, Sandra Fuchs, Jozée Sarrazin, Erin E. Easton, Alexandra Anh-Thu Weber

**Affiliations:** Department of Oceanography, University of Cape Town, Rondebosch 7701, Cape Town, South Africa; University of Brest- UMR 6539 CNRS/UBO/IRD/Ifremer, LEMAR – IUEM, Plouzané, France; Univ Brest, CNRS, Ifremer, UMR6197 BEEP, F-29280 Plouzané, France; Department of Zoology, University of Oxford, Oxford, United Kingdom; Nekton Foundation, Oxford, United Kingdom; School of Earth, Environmental, and Marine Sciences, University of Texas Rio Grande Valley, Brownsville, Texas, USA; Marine Invertebrates, Museums Victoria, Melbourne, Victoria, Australia; Department of Aquatic Ecology, Eawag, Swiss Federal Institute of Aquatic Science and Technology, Dübendorf, Switzerland

**Keywords:** international conference, inclusivity, opinion survey, pandemic, carbon footprint, meeting organization

## Abstract

Hybrid conferences are in-person events that are also accessible online. This type of meeting format was rare pre-COVID-19 but started to become more common recently given the asynchronous global progression of the pandemic and the uneven access and distribution of vaccines that led to a large proportion of participants being unable to attend international conferences in person. Here we report the organization of a middle-sized (581 participants: 159 onsite, 422 online) international hybrid conference that took place in France in September 2021. We highlight particular organizational challenges inherent to this relatively new type of meeting format. Furthermore, we surveyed both in-person and online participants to better understand their conference experience and to propose improvements based on the feedback received. Finally, we compare the advantages and disadvantages of three types of conferences (onsite-only, online-only and hybrid) and suggest that hybrid events should be favored in the future because they offer the most flexibility to participants. We conclude by proposing suggestions and ways forward to maximize accessibility and inclusivity of hybrid conferences.

## 1 Introduction

Scientific conferences are essential components of researchers’ lives, allowing them to stay up to date with the latest research trends while disseminating their work to the scientific community. These events are essential for networking and developing collaborations, especially for early-career researchers (ECRs; students and pre-tenure postgraduates) who use meetings as an opportunity to plan their next career step (Oester et al., 2017). Because of the COVID-19 pandemic and the resulting travel bans and restrictions, many in-person meetings since March 2020 were canceled, rescheduled, or changed to an online format, allowing scientists to present their research and interact with members of their respective communities virtually (Stefanoudis et al., 2021).

Online-only meetings have a number of advantages, for instance: (i) enhanced accessibility by allowing attendance during periods of fieldwork or teaching (Bartlett et al., 2021), (ii) reduced carbon footprint (Burtscher et al., 2020; Tao et al., 2021), and (iii) lower participation costs with potentially reduced registration fees and no travel and accommodation costs. These advantages have greatly improved inclusivity for researchers and students from developing countries and with limited financial means (Chou and Camerlink, 2021; Wu et al., 2022) and were the reasons why a large online international conference on photonics was held just before the COVID-19 pandemic (Reshef et al., 2020). Thanks to these advantages, many online conferences showed higher registration rates compared to previous in-person meetings (e.g., Castelvecchi, 2020; Stefanoudis et al., 2021). Online conferences, however, have a number of drawbacks, including: (i) fewer interactions among participants, especially if presentations are pre-recorded (Roos et al., 2020); (ii) increased fatigue after hours on screen (Bennett et al., 2021); (iii) fewer possibilities for spontaneous discussions and meetings (Roos et al., 2020); and (iv) technical issues during live talks resulting in schedule delays (Archibald et al., 2019).

Hybrid meetings, which have in-person attendance with a possibility to attend online, represent a promising solution that could address some of the shortcomings inherent of in-person or online-only meeting formats. There have been calls for adopting a hybrid format after all COVID-19 travel restrictions have been lifted (Joo, 2021), and there seems to be an interest amongst the scientific community for that format (Stefanoudis et al., 2021). However, due to the novelty of the hybrid format, conference organizers have to be creative to organize a successful event in which both in-person and online attendees are satisfied. So far, studies on hybrid meetings are scarce and focus on organizational and logistical aspects without accounting for the participant experience (Fulcher et al., 2020; Weiniger and Matot, 2021).

Here, we present information on the logistics of a recent international meeting, the 16th Deep-Sea Biology Symposium (16DSBS), a 5-day, medium-sized (581 attendees) hybrid conference that took place in Brest (France) in September 2021. We then compare the hybrid format to the in-person and online meeting formats in terms of costs and widening access. Finally, we report the participants’ experience using an online questionnaire to identify what worked well and less so. Based on those experiences we make some recommendations on how future organizers can improve the hybrid meeting experience.

## 2.1 Hybrid meeting logistics

### 2.1 Pre-meeting considerations

An important starting point for the organizing committee is defining the concept of the hybrid event, i.e., defining to what extent the online attendees participate in the conference. Can they be presenters, or do they only attend the conference? What is the expected level of interaction between and among onsite and online attendees? While informal interactions tend to form naturally among onsite attendees during coffee breaks and meals, these interactions are lacking for online attendees who usually need a screen break during these times. Hence, if the organizers wish that online participants interact among each other and with onsite participants, they have to organize special events to do so.

#### 2.1.1 How to choose a venue for onsite attendance?

The onsite organization for in-person attendance is analogous to a traditional in-person conference, and we thus focus mainly on the organizational aspects specifically related to the hybrid aspect. A major component of these events is that presentations should be recorded and live broadcasted, so infrastructure to support this component is essential at the selected venue. The required infrastructure can be: (i) provided by the venue (built-in cameras and sound system; personnel from the venue handling the retransmission); (ii) outsourced to an external company (an extra room is needed for the filming crew); and (iii) a static temporary camera installed / using the built-in cameras of laptops (with members of the local organizing committee (LOC) handling the retransmission, for instance via zoom). A combination of these options is also possible.

#### 2.1.2 Which platform(s) to choose for online attendance and communication?

The choice of an appropriate online platform for a hybrid conference is crucial because it should ideally (i) provide easy access to the online content of the conference (live talks; on-demand talks; posters) and (ii) aim to enhance all types of exchange and communication among onsite and online participants (e.g., live chat).

For pre-meeting communications, emails and a dedicated website are usually the best solution. However, they may not be the best way to communicate with online and onsite attendees during the conference. Rapid messaging through a dedicated mobile application for the conference, or via online platforms (e.g., Slack), is an efficient way to communicate important information rapidly. Important aspects to take into account are: (i) making sure that all participants have access to these messaging platforms and (ii) providing enough time to participants to become familiar with these platforms.

#### 2.1.3 Which format to choose for presentations?

##### Talks

While presenting live is the norm for onsite presenters, it is more challenging for online presenters. For online speakers, giving a live talk has a number of advantages, such as more interactions and the possibility to answer questions live. However, it also has a number of drawbacks, such that time slots for talks will inevitably not be suitable for the time zones of all participants, and live online talks are more prone to technical issues that can result in delays. Organizers should decide which option(s) they want to give online presenters, such as (i) presenting live and answering questions live, sending a pre-recorded talk but answering questions live and (iii) sending a pre-recorded talk and not being present for questions (e.g., if time zones are incompatible). Offering all three options is the most flexible for online speakers, however this flexibility entails more expense, organization, and risks of delay.

##### Posters

In-person poster sessions are not different from a classical onsite-only conference. However, in-person and online posters should be available to view on the conference platform. Ideally, a chat box next to each poster should be accessible for questions and answers, and a live online poster session should be organized to allow for live interactions with online presenters.

#### 2.1.4 What additional considerations does the hybrid format entail from an organizational perspective?

Organizing a hybrid conference entails the usual logistics required for an in-person-only and an online-only conference, but there is additional work for the LOC that is inherent to the hybrid format.

##### More communication, flexibility and file handling

Clear communication with participants is essential and common to all conferences but the hybrid format adds complexity due to several types of participation. For instance, any registration change (e.g., onsite to online, or vice-versa) has to be followed by updates in internal databases, mailing lists and the program. Communication efforts also increase because customized instructions have to be provided to online and onsite participants and presenters. Furthermore, a considerable amount of work has to be done ahead of the conference to receive and organize all presentation files (e.g., pre-recorded talks and posters).

##### More complexity to design the program

The hybrid format typically implies a larger participation, compared to in-person conferences, which can result in more requests for talks and thus competition for the available time slots. Ideally, the talk schedule should be organized according to the time zone of the online speakers. However, this consideration is not always compatible with the scientific sessions and venue hours of operation. To avoid organizing a two-tier conference with onsite participants getting much more interactions than online participants, the LOC should organize online-only events beyond talks and posters to enhance interactions among online participants and between onsite and online participants.

##### More support personnel

The above-mentioned tasks require increased administration pre-conference workload for the LOC. Furthermore, during the conference, additional chair and co-chair persons are needed to facilitate question-and-answer sessions from the onsite and online audience (passing on microphones; checking the chat box). To increase inclusivity, chairs can be online participants, however, an onsite co-chair would also be needed. Finally, members of the LOC are also required to moderate online-only events and respond to the requests of online attendees.

### 2.2 Case study

The Deep-Sea Biology Symposium is an international in-person conference organized every three years by the Deep-Sea Biology Society (DSBS) and a LOC. For reasons related to the global COVID-19 pandemic, the French Research Institute for Exploitation of the Sea (Ifremer) was asked to replace the planned LOC for the 16th Deep-Sea Biology Symposium (16DSBS) approximately a year before the event took place. The symposium was held 12-17 September 2021 in Brest, France, at the Aquarium Océanopolis. The conference consisted of two parallel sessions divided into two rooms: a room which had built-in cameras suitable for broadcasting managed by the personnel venue and a second room in which an external company was hired to organize the live broadcasting. This company also set up the streaming website on which all live and on-demand talks could be watched up to two weeks post-conference. The team of the conference venue was formed by two people in the control room and one sound engineer; while the external company consisted of a crew of five people: two people in the control room, one sound engineer and two cameramen.

In terms of scientific content, the 16DSBS contained 214 contributed talks (64% acceptance rate) and 170 posters over five days (Fig. 1; File S1). To enhance their visibility, poster presenters were asked to provide a 2-min video pitch of their poster, in addition to a PDF and/or a printed version of their poster, depending on their attendance type. In addition, in order to maximize the participation of online attendees, we organized a total of 11 online-only events across different time zones. These were: an early career researcher/student mixer; five zoom lunches with keynote speakers of the day; a round table on decolonizing deep-sea science; a 3-hour poster session; an online Gala dinner with social activities, and the annual general meeting of the Deep-Sea Biology Society. The conference was attended by 581 participants, with approximately three quarters of them attending online (Fig. 1). Finally, both onsite and online participants could present either talks or posters (the talk selection process did not take attendance type into account); live or pre-recorded for online participants.

**Figure 1:**
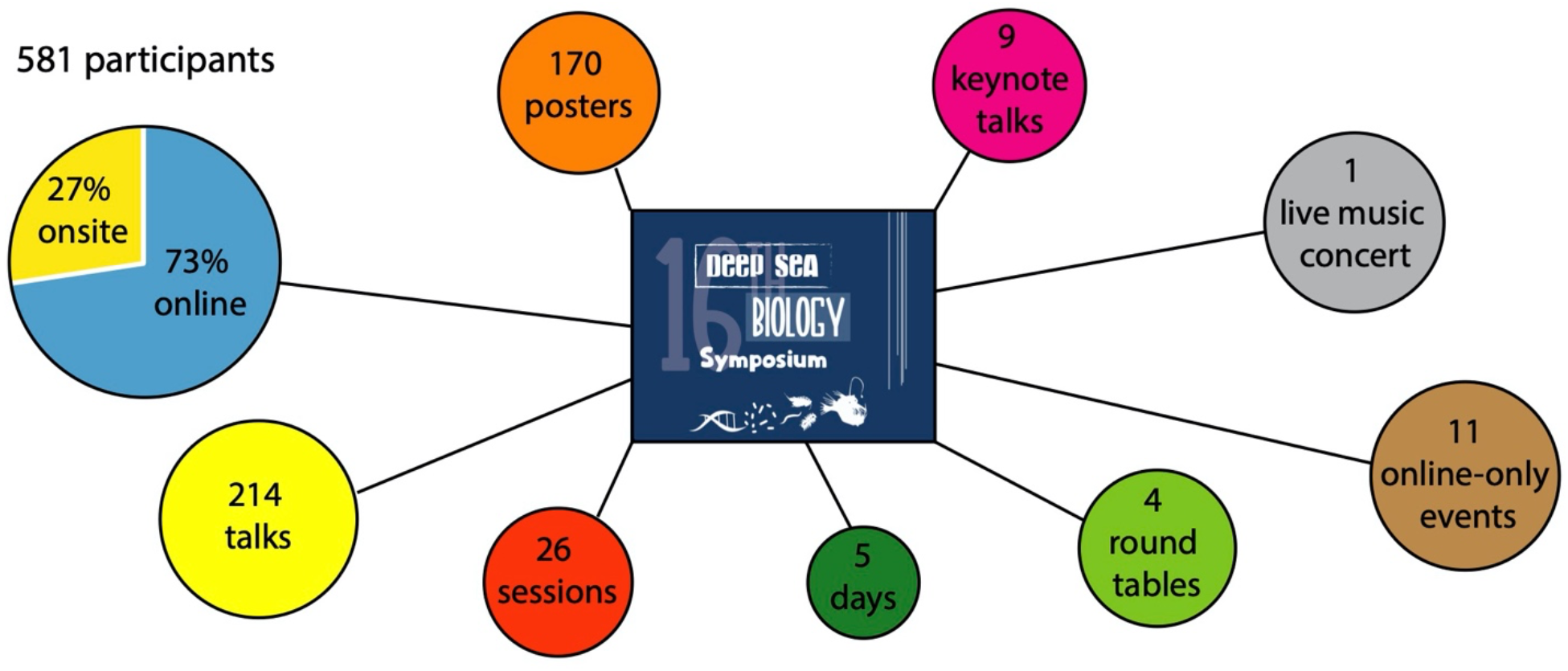
Summary of attendance and content of the hybrid conference 16DSBS. The 26 scientific sessions were presented in two parallel sessions.

#### 2.2.1 Pre-meeting organization

A conference website with all pre-conference information was hosted on servers of Ifremer (Table 1). A dedicated email address was created including relevant mailing lists to address different participants (e.g., all attendees; onsite only-attendees; online-only attendees; all presenters (talks & posters)). Online attendees were offered the choice to (i) present live and answer questions live (ii) send a pre-recorded talk but answer questions live, or (iii) send a pre-recorded talk and not be present for questions (e.g., if time zones were incompatible). Online presenters were asked to send a pre-recorded version of their presentation to be used as a backup. We aimed to obtain a maximum of live talks, and we thus adjusted the talk schedule according to the time zone of online speakers. However, it was not always possible due to each talk being scheduled within its relevant scientific session of which there were 26.

**Table 1:** Summary of online platforms used for 16DSBS and their purpose.

| Description | Access | Aim | Cost |
| --- | --- | --- | --- |
| Conference website | Open | General information; registration; abstract submission | Supported by organizing institution Ifremer – not in conference budget |
| Conference email address | Open | Pre-conference communications | Supported by organizing institution Ifremer – not in conference budget |
| Streaming channel/website | Password-protected | Live and pre-recorded talks; talks on replay; live chat from online audience for questions to speakers | 46% of total budget; see Fig. 2 |
| Private page on conference website | Password-protected | Access to posters in pdf format; links to short video pitches of the posters | Supported by organizing institution Ifremer – not in conference budget |
| Private YouTube Channel | Private link | Access to short poster videos | Free |
| Zoom | Password-protected | Live talks from online speakers; Online poster session; online-only events | Supported by the DSBS – not in conference budget |
| Slack | Invitation via email | Communication before and during the conference; sharing of different passwords; asking non-live questions | Free |
| Twitter conference account | Open | Communication and public engagement before and during the conference | Free |
| Gather Town | Open | Networking and social events | Free |

#### 2.2.2. Online access to the conference

At the time when the 16DSBS was organized, there was no single online platform available to host all online content of a hybrid conference. Furthermore, outsourcing the development of such a platform was out of financial reach for the society-based 16DSBS. Hence, a streaming channel including (i) live talks, (ii) chat box for live questions from the online audience, and (iiii) on-demand talks was developed by the external company hired for the live filming and broadcasting (https://16dsbs.attwm.fr). For other online content (e.g., access to online posters; online-only events; etc), we relied on free platforms or platforms whose costs were covered by the hosting institution Ifremer and the Deep-Sea Biology Society. Overall, this resulted in a large number of different platforms (Table 1).

#### 2.2.3 Budget

For the hybrid 16DSBS, the total budget was slightly lower than an estimated budget for a same-sized onsite-only conference (Fig. 2). Details of the different budgets are provided in File S2. We refer to the costs provided by Stefanoudis et al., (2021) for a budget for an online-only event. The budget for an onsite-only event was estimated using the onsite costs of the 16DSBS projecting the costs associated with 200 people expected to attend onsite to 581 attendees, which was the total number of 16DSBS online and onsite attendees.

**Figure 2:**
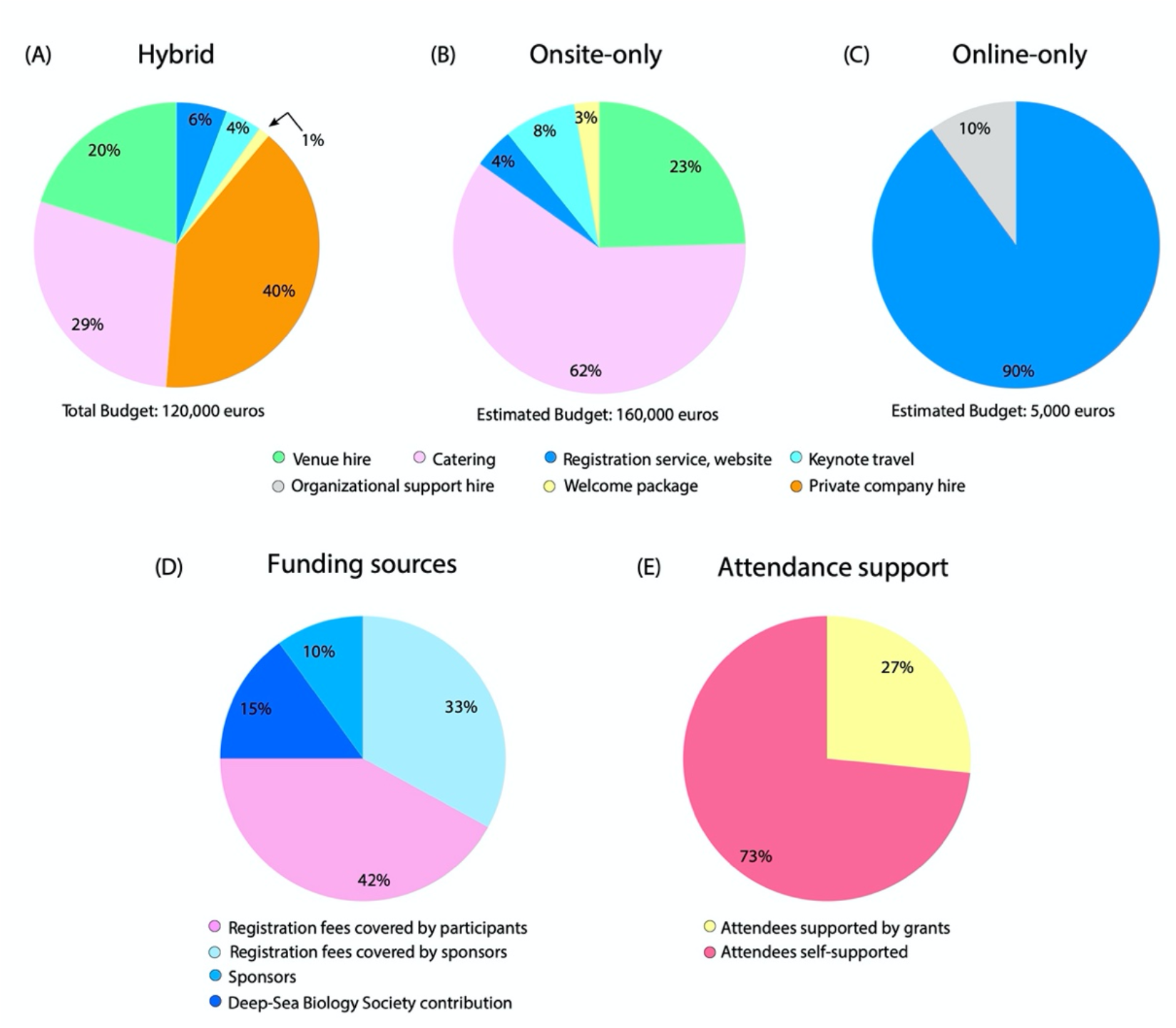
Relative contribution (%) of the budget from the three types of conference: (A) hybrid, (B) online only and (C) onsite only. The budget in (A) reflects the total costs of the hybrid 16DSBS. Budget estimates are based on a conference with 581 participants: (B) if it was hosted onsite-only at the Océanopolis Aquarium and (C) if it was organized exclusively online following the budget of eDSBS, an online-meeting (Stefanoudis et al., 2021). Organizational support hire includes costs associated to hire staff member(s) dedicated to the event organization. (D) General proportion of the funding sources for 16DSBS, including the amount of the registration costs covered by sponsors, in particular the DSBS and the International SeaBed Authority. (E) Relative proportion of attendees who were supported by travel grants offered by the DSBS together with the International SeaBed Authority. The total number of participants was 581.

Compared to this estimated budget, the 16DSBS catering and food service fees were reduced and audio-visual costs were higher. Specifically, a significant part of the 16DSBS budget was dedicated to the hire of a professional company (5 people) that (i) organized the filming of in-person talks for one session, (ii) organized the live broadcasting, (iii) ran pre-tests with online speakers, (iv) set up the streaming website for live talks and on which recorded talks were available on demand for two weeks after the conference, and (v) uploaded the recorded talks at the end of each day. This service could not have been accomplished by the LOC itself. To minimize registration fees for attendees, the LOC decided to use other platforms for the other events and presentations (Table 1); however, this cost-saving measure increased the complexity of navigating among platforms for online participants. Another relevant cost is represented by hiring dedicated staff member(s) for the organization of the conference. In our case, two people were specifically hired for one year to organize the event, however this cost was supported by Ifremer, and thus did not affect the final budget.

While the hybrid 16DSBS and the estimated onsite-only conference budgets are similar, the estimated budget of an online-only conference of a similar size is considerably reduced (Fig. 2). Indeed, expenses for virtual conferences exclude most in-person conference costs except for administration and registration and website platforms. Nevertheless, as for hybrid conferences, additional costs are incurred for virtual platforms to host the conference and cloud storage costs to make presentations available for a designated time (Fig. 2) (Stefanoudis et al., 2021).

For the 16DSBS, 42% of the costs were covered by the registration fees of participants. The remaining funding was acquired by the LOC through sponsoring or from contributions of the Deep-Sea Biology Society (DSBS) (Fig. 2d). Financial support from sponsors and the DSBS was provided either as direct payment to the LOC (25%) or in the form of travel/registration grants to attendees (33%).

The 16DSBS registration fees for online-only attendance were lower than onsite-only, and rates for student/researcher from developed and developing countries were not differentiated (Table 2). Registration costs for online attendance and the holding of an in-person event raised a debate within the deep-sea community for a few months prior to the event. Notably, critics reported (i) the inaccessibility for some prospective attendees to afford such costs and (ii) the inability for many to participate onsite due to travel restrictions associated with the COVID-19 pandemic. While the LOC acknowledges that it may have lacked transparency during the organizational phase, it uses the present article to provide some clarity and perspective. First, it should be noted that overall budgets for online-only and hybrid conferences are very different (Fig. 2A-C), which is inevitably transferred to registration costs to some extent. Second, the 16DSBS online registration costs are within or below the range of other hybrid conferences held in 2021 and 2022 (Table 2). And third, about a quarter of attendees (mainly ECRs and researchers from developing countries) were supported by travel/registration grants (Fig. 2E).

**Table 2:** Non-exhaustive examples of registration fees for 2021-2022 hybrid conferences. Ranges include all rates from the highest discounts, generally for students, society members, and low-income countries, to the maximum costs for onsite registration. All registration costs have been converted to euros to simplify comparisons (the original prices are in brackets). Conference information last accessed on 22 February 2022.

| Name | Dates | Onsite registration | Online registration | Online speakers | website |
| --- | --- | --- | --- | --- | --- |
| 16th Deep-Sea Biology Symposium | 12.09.21-17.09.21 | 380-600 EUR | 100-375 EUR | yes; interspersed (live or pre-recorded) | https://wwz.ifremer.fr/16dsbs/ |
| Annual Meeting of the Association for Information Science and Technology | 30.10.21-02.11.21 | 311-745 EUR (350-850 USD) | 22-439 EUR (25-495 USD) | yes (live or pre-recorded) | https://www.asist.org/am21/ |
| International Conference on Biodiversity, Ecology and Conservation of Marine Ecosystems | 03.01.22-07.01.22 | 139-276 EUR (161-321 USD) | 70-139 EUR (81-161 USD) | yes (format not specified) | https://www.become2022.com |
| 2022 Conference on General Education, Pedagogy, and Assessment | 10.02.22-12.02.22 | 386-582 EUR (75-675 USD) | 110-241 EUR (50-275 USD)* | yes; single day (format not specified) | https://www.aacu.org/events/2022-conference-general-education-pedagogy-and-assessment |
| 2022 Conference on Diversity, Equity, and Student Success | 17.03.22-19.03.22 | 386-582 EUR (75-675 USD) | 110-241 EUR (50-275 USD)* | yes; single day (format not specified) | https://www.aacu.org/events/2022-conference-diversity-equity-and-student-success |
| American Educational Research Association | 21.04.22-26.04.22 | 83-846 EUR (95-590 USD) | 57-605 EUR (65-485 USD) | yes; dedicated sessions (format not specified) | https://www.aera.net/Events-Meetings/Annual-Meeting |
| 2022 World Aquaculture and Fisheries Conference | 18.05.22-19.05.22 | 648-911 EUR (739-1039 USD) | 385-560 EUR (439-639 USD) | yes (live or pre-recorded) | https://www.worldaquacultureconference.com/ |
| World Biodiversity Forum 2022 | 26.06.22-01.07.22 | 312-768 EUR (325-800 CHF) | 144-240 EUR (150-250 CHF) | yes (format not specified) | https://www.worldbiodiversityforum.org |
| 16<sup>th</sup> World Congress on Bioethics | 20.07.22-22.07.22 | 210-577 EUR (225-620 CHF) | 210-577 EUR (225-620 CHF) | no | https://iab2022.org/ |
| 24<sup>th</sup> Biennial Conference on the Biology of Marine Mammals | 01.08.22-05.08.22 | 131-701 EUR (150-800 USD) | 88-351 EUR (100-400 USD) | yes; dedicated sessions (pre-recorded) | https://www.smmconference.org/ |

\*Online includes a single day of virtual conference and live streaming of plenary on the remaining days.

## 3. Participants’ perspective

### 3.1 Participation in comparison with previous meetings

Comparisons with previous deep-sea-biology-themed meetings indicate a marked increase in participation, 49% against an in-person meeting in the USA (15DSBS), 343% against an in-person meeting in Colombia (ISDSC7, which had a narrower scientific focus on deep corals), and 65% against an online meeting (eDSBS, Table 3). The proportion of ECRs also increased (55% of all participants) in comparison to in-person meetings (25-36%) but decreased to the online-only meeting (65%) that prioritized ECR presentations (Stefanoudis et al., 2021) (Table 3). This enhanced ECR participation also translated into more presentations delivered by ECRs (57%) compared to 23% (15DSBS), 42% (ISDSC7), and 79% (ECR-focused eDSBS).

**Table 3.**
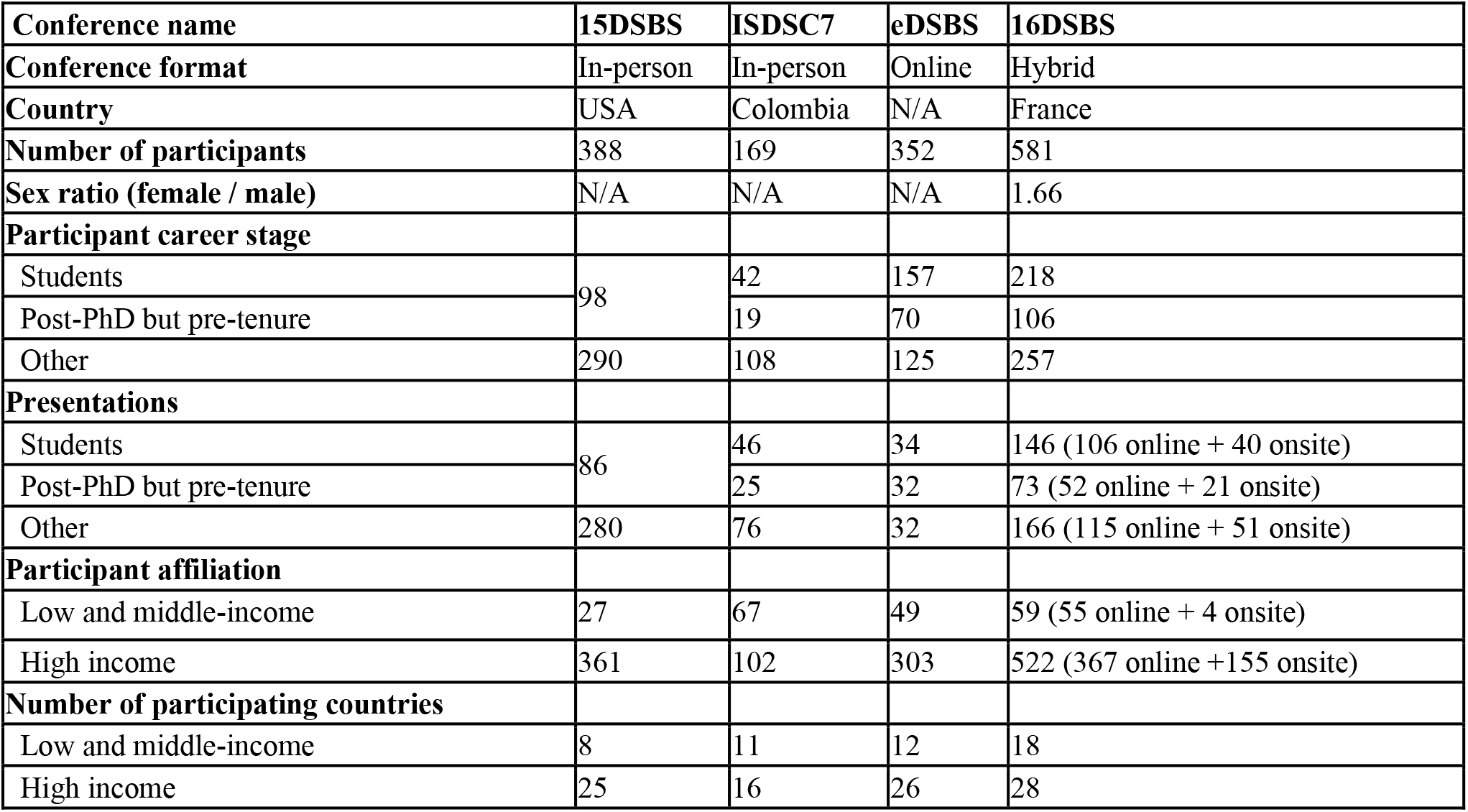
Comparison between in-person, online and hybrid deep-sea biology meetings in terms of demographic composition by sex, career stage and country of institutional affiliation. Sex ratio estimates excluded non-binary, or those that chose not to disclose sex. Students include PhD candidates too, while tenure includes any equivalent permanent position. For 15DSBS, no separation was made between students and post-PhD but pre-tenure. Number of participating countries was identified from participants’ institutional affiliations. Country categories based on the 2021 classification by the World Bank (last accessed on 16 February 2022). N/A = Not applicable.

Furthermore, the proportion of participants from low and middle-income countries represented at the hybrid 16DSBS was 11%, which was lower than the eDSBS (14%) and ISDSC7 (40%), but higher than the 15DSBS (7%). It should however be noted that in terms of total low- and middle-income participants, the 16DSBS was the second highest following the ISDSC7 in Colombia (Table 3). Overall, there is poor representation of low- and middle-income researchers in the field of deep-sea biology (Costa et al., 2020), and although the hybrid format can aid participation of those researchers, holding an in-person meeting or the in-person aspect of a hybrid meeting in a low- to middle-income nation can be much more effective in widening participation.

### 3.2 Questionnaire for participants

To gather impressions and feedback from participants, we organized an online survey focusing on the16DSBS content and organization. The full questionnaire and replies are available in the supplementary File S3. A total of 164 participants replied (28% of total participants), 104 online (25% of online) and 60 onsite (38% of onsite) participants.

#### 3.2.1 Meeting format and technical considerations

From a technical perspective, both online and onsite participants enjoyed the live conference experience (72-92%, Q17-20), with all platforms utilized (i.e., to allow online participation, online Q&A, and accessing of live and recorded presentations) deemed as sufficient and easy-to-use (57-74%, Q21,23,30-31). However, a sizable proportion found that the number of platforms used was too high (38%, Q32) and suggestions for future usage of fewer and more all-encompassing platforms were made (See Supporting information). Moreover, most agreed with the number of talks allocated per day and the overall duration of the conference (75%, Q35) and did not support a future third parallel session to accommodate more talks (66%, Q41).

The majority of participants regarded live talks as an integral part of the conference (79%, Q11) as it enhances interactions (60%, Q36). The option of pre-recorded talks to cater for those with technical issues or time zones differences was considered essential (84%, Q12). The on-demand feature was overwhelmingly well-received (92%, Q13) with most indicating they viewed content post-conference (79%, Q14). However, opinion was split on if the 2-week post-conference availability of that feature was adequate (Q15), with some suggestions to increase the duration to a month or more in the future (See Supporting information).

Most agreed with the format of online posters (64%, Q42) and found the additional short-video pitch useful (66%, Q43), although it should not substitute the pdf file of the poster (70%, Q46). There were mixed feelings on the duration of the poster sessions, with more satisfaction for onsite (55%, Q44) compared to online (43%, Q45), although it is not clear from the questionnaire and comments received if the session should have been shorter or longer. Based on comments received, it became apparent that future hybrid meetings should aim to better link online and onsite poster sessions, perhaps by including Q&A sessions with presenters, either live (69%, Q47) or online (70%, Q48).

#### 3.2.2. Networking

In terms of networking more than two thirds indicated that they were able to connect with other researchers (70%, Q70), although the number of questions they received compared with past in-person or online meetings was less for 48% and 54% of onsite and online participants, respectively (Q72-73). The latter finding is interesting, and is probably best explained by the fact that the majority of online (54%) and onsite (69%) participants did not interact with the other group of participants (Q74-75), with only 44% of all participants engaging with both groups (Q71), thus limiting the number of potential interactions per participant.

Several online social events were organized to enhance the online conference experience, most of which were generally well-liked, including the early-career focused scientific illustration workshop (64%, Q58), the lunch-time social gatherings events with the respective keynote speaker of the day, (80%, Q59), and the online Gala activities (88%, Q67). However, comments indicated participation in these events was limited by time-zone conflicts and from onsite attendees, with only 22% of onsite attendees indicating that they participated in several online social events (Q62).

#### 3.2.3. Overall experience and moving forward

The vast majority of participants agreed or strongly agreed that the conference was an enjoyable experience (88%, Q2), inclusive (72%, Q3) and of high scientific quality (72%, Q4). Online and onsite attendees experienced the event slightly differently, with the former finding it more difficult to concentrate (39% vs. 22%, Q5-6) and dedicate time (53% vs. 18%, Q7-8) for this meeting compared to past in-person meetings. Time zone conflicts and work duties (teaching, lab work) were some of the reasons evoked by online participants. There were mixed feelings about the amount of registration fees (Q9), although approximately half agreed that awards from the Deep-Sea Biology Society were sufficient to cover registration and attendance fees for those in need (52% agreed vs. 10% disagreed, Q10).

Moving forward, the overwhelming majority of participants indicated that they want future Society-sponsored meetings to be hybrid (79%, Q81), with considerably less appetite for future in-person only or online-only events (11% and 21%, respectively, Q82-83). Finally, additional small online events, including webinars, lectures series and journal clubs, to be held between symposia were largely favored as well (79%, Q84).

## 4. How to organize a hybrid conference?

### 4.1 Summary

This paper highlights what the organization of a medium-sized hybrid international conference entailed in 2021. As organizers, we report our experience and gathered feedback from both types of delegates to highlight successes and possible ways of improvement. Below we highlight key relevant points that should be accounted for and possible solutions to improve the organization of such events in the future.

### 4.2 Advantages and disadvantages of the hybrid format

We summarized the pros and cons of the three types of existing meetings in Table 4 (onsite-only, online-only, hybrid). Overall, we believe that hybrid meetings are better than onsite-only or online-only meetings for participants because they offer more flexibility to delegates. Indeed, for those who can travel, they provide the much-needed in-person interactions offered by onsite meetings while offering the possibility to attend online for researchers with limited financial means, other commitments (e.g., work or care duties) or who do not wish to travel for environmental reasons. Indeed, hybrid meetings have overall lower carbon footprints than similar-sized onsite-only conferences. However, there are two main downsides to hybrid meetings: (i) they are more complex to organize (see section 2.1.4), which can lead to (ii) generally more expensive meetings for online participants (but see section 4.3).

**Table 4.** Summary of advantages and disadvantages of onsite-only, online-only and hybrid meetings for participants.

| Onsite-only |  | Online-only |  | Hybrid |  |
| --- | --- | --- | --- | --- | --- |
| Advantages | Disadvantages | Advantages | Disadvantages | Advantages | Disadvantages |
| Extensive opportunities for networking and social interactions | High registration and travel costs limits accessibility | Reduced registration costs and absence of travel / accommodation expenses enhances accessibility | Limited opportunities for networking and social interactions | More flexibility for delegates implies an overall higher participation | Overall higher organization costs compared to online-only meeting may lead to higher registration fees for online participants compared to online-only conferences |
| Visiting a new city or country; learn about a new culture | High carbon footprint due to travel | Low carbon footprint | Screen fatigue | Extensive opportunities for networking and social interactions (onsite participants) | Online participants may feel excluded from networking and social activities |
|  | Incompatibilities with other commitments (e.g., teaching, fieldwork, lab work, personal life) | Recorded presentations can be available post-conference for a given time | Potential time zone issues | Recorded talks can be available after the conference for a given time for all participants | Increased workload for the organizing committee (e.g., general organization; program scheduling; communication with online and onsite attendees) |
|  | Typically, no recording of presentations | Limited schedule delays if pre-recorded | Without subtitles, might be more | Reduced international travel for online |  |
|  |  | presentations are<br>broadcasted | exclusive for people<br>with impaired hearing | participants: decreased<br>carbon footprint |  |
|  |  | Adding subtitles to pre-recorded talks may aid<br>non-native speakers<br>and/or people with<br>impaired hearing |  | Adding subtitles to pre-recorded talks from<br>online speakers may aid<br>non-native speakers<br>and/or people with<br>hearing impairments |  |

### 4.3 Recommendations for future hybrid meetings

1. **Define clearly the extent of online participation** As mentioned in the introduction, a substantial part of hybrid meeting complexity and increased costs is linked to the extent of participation of online attendees. If they can only attend the conference without presenting, the complexity decreases drastically, however it disadvantages individuals not able to travel. Furthermore, if they can present, offering them the choice to present live or ask them to send a pre-recorded talk (or present both options) adds another layer of complexity. Finally, organizers can also choose to what extent they wish to organize extra online-only events beyond talks and posters to maximize interactions among online attendees. We believe it is fairer that online attendees can also present their research in the way it suits them best, and that they have a number of opportunities to network. Indeed, scientific conferences are not only meant to present one’s research, but also interact with the members of their own community. However, the more options there are for online attendees, the more work there is for the organizing committee, which may translate in higher registration costs for everyone, especially online attendees. For each hybrid event, there is a fine balance to find between offering the best experience for all types of attendees, and keeping registration costs low without overwhelming the organizing committee.
2. **Maximize inclusivity** We believe that the main aim behind organizing hybrid conferences is to broaden the participation of scientists by offering them flexibility for the attendance type. Hence, providing an inclusive conference is likely a goal of each organizing committee. While registration costs of the 16DSBS were similar to or lower than those of other hybrid conferences (Table 2), we acknowledge that registration costs for students or researchers from low- and middle-income countries could have been differentiated, and thus even lower. Here are some propositions to maximize inclusivity of a hybrid event: (i) reach out to numerous sponsors to lower registrations fees; (ii) differentiate registration fees by attendance type (online; onsite), career stage (student; post-doc; established researcher), nation income category (low, medium and high-income country), if it is a society meeting (member; non-member), time of registration (early bird; regular; late registration), (iii) provide additional travel and registration awards; (iv) aim to have the meeting organized by developing nations; and (v) reach out to the scientific community before the event (e.g., via online surveys) to better understand individual needs and challenges. Finally, as participants are not necessarily aware of the additional logistical requirements needed for hybrid conferences, we suggest publishing a cost breakdown along with registration fees on the conference website. Hence, potential higher costs of hybrid events in comparison with online-only events are better justified.
3. **Simplify (online) access and communication** We received a recurrent negative feedback from online attendees: there were too many platforms to access the conference and interact with other online attendees and their use was too complicated (Table 1). We acknowledge this issue, however, at the time when we organized the 16DSBS, there was no single platform available for this kind of event. Furthermore, the set-up of a dedicated platform for the purpose of this conference by an external company would have increased costs. In addition, efficient communication to all participants before and during the conference was not optimal. A large number of emails were sent before the conference. During the conference, we attempted to use Slack to communicate rapidly with all participants, however, it was mainly online participants who used it, and not everyone did use it (there was some reticence from first-time users). We thus recommend future organizers to aim for a single platform to access live and on-demand talks, posters, ask questions to speakers, and more generally interact with other online attendees via chats or videoconferences, as well as to receive information from and communicate with the organizing committee for potential issues. Ideally, we suggest that a combination of a dedicated website for viewing and a linked mobile app for rapid communication would be the best option. Nevertheless, we realize this centralization is a difficult endeavor, and hope that in the future such platforms will exist or their set-up by external companies will be more affordable.
4. **Maximize interaction opportunities between online and onsite attendees** Finally, while onsite and online participants had equal access to scientific talks, we noticed that for the remaining scientific activities (e.g., poster sessions; online-only events) the two types of delegates were not really interacting with each other. For instance, onsite participants appeared to prefer getting some rest or interacting with onsite colleagues rather than participating in online-only events. Furthermore, online participants did not have an easy way to interact with onsite participants if the latter were not using Slack. We realize that there is probably no way to fully overcome this issue, however, we believe that organizers should aim at minimizing this problem. For instance, developing a mobile app that all participants would have to download will likely make communication and networking among all attendees easier (e.g., II Joint Congress on Evolutionary Biology, Montpellier, 2018).

### 4.4 Conclusion

Despite some organizational challenges, we advocate to keep organizing hybrid conferences beyond the COVID-19 pandemic. Indeed, they allow for a wider participation by giving more flexibility to participants to choose an attendance type that suits their needs best. Furthermore, online-only conferences cannot fully replace in-person formal and informal interactions. Although hybrid events require additional work and are currently more expensive in comparison to online-only events for online participants, we think that with early planning, sufficient sponsors, and technological advances, hybrid events represent the most inclusive way to hold international conferences.

Furthermore, hybrid conferences have lower carbon footprint compared to onsite-only conferences, hence they offer an interesting opportunity to combine scientific networking with environmentally-friendly decisions (Glausiusz, 2021). For instance, students and researchers could decide to attend conferences in-person whose locality is reachable by train, while attending online conferences taking place on other continents.

We would like to emphasize that having an online option for a conference should not become an excuse for institutions and funding sources not to fund students and researchers to attend the conference onsite anymore. In-person networking is an essential part of a researcher’s work to develop collaborations, especially for ECRs who can find their next career step during these events. In addition, we would encourage the in-person element of international hybrid meetings to take place in low- and middle-income nations as it enhances diverse participation or to change continents to benefit all geographies equally. As hybrid conferences become more common, their organization may become more straightforward. This article reports the organization of one of the first hybrid conferences, and we hope that our experience will be valuable to the organizers of future hybrid events.

## Conflict of Interest

The authors declare that the research was conducted in the absence of any commercial or financial relationships that could be construed as a potential conflict of interest.

## Author Contributions

DZ, AWH, AATW, EP, MK, SF, JZ, MM and LM were members of the 16DSBS local organizing committee. AATW initiated and led manuscript preparation and writing. All authors contributed to the survey design. PS, LM and SF analyzed the survey data and drafted the corresponding sections. EP and EE drafted the budget section. AATW, EP and PS drafted the remaining sections of the manuscript, with feedback from all co-authors.

## Funding

This work and EP were supported by the ISblue project, Interdisciplinary graduate school for the blue planet (ANR-17-EURE-0015) and co-funded by a grant from the French government under the program “Investissements d’Avenir”. AATW was funded by a Marie Sklodowska-Curie Global Fellowship awarded by the European Union’s Horizon 2020 Research and Innovation Program (Grant No. 797326).

## Acknowledgments

We are grateful to all members of the 16DSBS local organizing committee (alphabetical order): Alizé Bouriat, Marie-Anne Cambon, Valérie Cueff-Gauchard, Titouan Doussin, Rémi Dulermo, Lucile Durand-Le Rolland, Nahia Etchegaray, Didier Flament, Vincent Georges, Anne Godefroy, Nadine Lanteri, Marjolaine Matabos, Lénaïck Menot, Eve-Julie Pernet, Garance Perrois, Justine Pittera, Florence Pradillon and Pierre-Marie Sarradin. We are also grateful to the following trustees of the Deep-Sea Biology Society (alphabetical order): Chong Chen, Malcolm Clark, Adrian Glover, Steven Haddock, Santiago Herrera, Raissa Hogan, Ilysa Iglesias, Rachel Jeffreys, Andrea Quattrini, Julia Sigwart, and Chris Yesson. We are also grateful to Amy Baco-Taylor, Giulia La Bianca, Mackenzie Gerringer and Guilherme Siqueira Toledo de Carvalho who organized the online Gala dinner and activities.

We would like to particularly thank the 164 respondents of the 16DSBS survey who gave their time to provide feedback on the conference. We believe that their opinion will be invaluable to improve future hybrid meetings. We are grateful to the employees of An Tour Tan (Tom Gonzalez and his team) and Océanopolis (Marie-Pierre Jacolot, Guy Bescond, Didier Harel, Laurent Dubouchet-Poncey) for logistical support. Finally, we would like to thank all conference sponsors (alphabetical order): the Campus Mondial de la Mer, the city of Brest, the department Finistère, the Deep-Sea Biology Society, the Gordon and Betty Moore Foundation, Ifremer, the International Seabed Authority and ISBlue.

## Supplementary Material

**File S1:** 16DSBS program overview

**File S2:** Detailed budget for same-sized: (i) 16DSBS hybrid conference, (ii) onsite-only conference (estimation for 581 people), (iii) online-only conference (estimation for 581 people).

**File S3:** 16DSBS post-conference questionnaire with responses.

## Data Availability Statement

The dataset analyzed for this study (16DSBS and participant replies) can be found in the supplementary material File S3.

## Notes

### Competing Interest Statement

The authors have declared no competing interest.

