## Supplementary File S1 for "Hybrid conferences: opportunities, challenges and ways forward"

### 16th DSBS

### Deep-Sea Biology Symposium

#### Brest, 12-17 September 2021

**Room Legend**

| Room |
|---|
| Auditorium |
| Pavillon Evènementiel |
| **Online only** |

| **Sunday 12/09** | |
|---|---|
| 16:30 | DSBS Registration |
| 18:00 | Icebreaking |
| 19:00 | **Online only**: **Combined ECR/Student Mixer** |

| **Monday 13/09** | | |
|---|---|---|
| 08:20 | **Welcome**, president address & local repres. | |
| 09:00 | **Keynote: Sabine Gollner** | |
| 09:35 | Special 1g/1h: **Deep sea mining** 6 talks | General 2: **Biodiversity** 6 talks |
| 11:05 | Coffee Break * 20 mins | |
| 11:25 | Special 1g/1h: **Deep sea mining** 7 talks | General 2: **Biodiversity** 7 talks |
| 13:10 | Lunch * 1.05 hours | |
| 13:30 | Lunch Time Social Event | |
| 14:15 | **Keynote: Amy Baco** | |
| 14:50 | Special 1g/1h: **Deep sea mining** 6 talks | General 2: **Biodiversity** 6 talks |
| 16:20 | Coffee Break * 20 mins | |
| 16:40 | General 1: **Conservation** 4 talks | Special 1b: **spatial planning** 4 talks |
| 18:00 | **Online only**: Student event: Creating impactful graphics and figures to showcase your science. DSBS | |
| 19:30 | **Online only**: Round table: Decolonizing deep-sea science. DSBS | |

| **Tuesday 14/09** | | |
|---|---|---|
| 08:30 | **Keynote: Moriaki Yasuhara** | |
| 09:05 | Special 1f: **Climate change** 4 talks & special 1c: **managing deep sea** 4 talks | Special 2a: **In honour of Craig Smith** 8 talks |
| 11:05 | Coffee Break * 20 mins | |
| 11:25 | Special 1e: **Pollutants** 4 talks & **Special: Communication** 3 talks | Special 2a: **In honour of Craig Smith** 7 talks |
| 13:10 | Lunch * 1.05 hours | |
| 13:30 | Lunch Time Social Event | |
| 14:15 | **Keynote: Kerry Sink** | |
| 14:50 | Special 2c: **NW Pacific** 6 talks | Special 2a: **In honour of Craig Smith** 6 talks |
| 16:20 | Coffee Break * 20 mins | |
| 16:40 | Poster session (between the 'Pavillon Polaire' and the 'Pavillon Tropical') & online | |

| **Wednesday 15/09** | | |
|---|---|---|
| 08:30 | **Keynote: Christian Tamburini** | |
| 09:05 | Special 2d: **Bioluminescence** 4 talks & Special 2e: **Microbiology** 4 talks | General 2: **Biodiversity** 8 talks |
| 11:05 | Coffee Break * 20 mins | |
| 11:25 | Special 2e: **Microbiology** 2 talks & Special 2f: **Microbiology** 5 talks | General 2: **Biodiversity** 7 talks |
| 13:10 | Lunch * 1.05 hours | |
| 13:30 | Lunch Time Social Event | |
| 14:15 | Social activities | |
| 20:00 | Gala Dinner | |
| 22.00 | **Virtual social event** | |

| **Thursday 16/09** | | |
|---|---|---|
| 08:30 | **Keynote: Johanna Weston** | |
| 09:05 | Special 4a: **Vision** 8 talks | Special 2b: **Hadal** 8 talks |
| 11:05 | Coffee Break * 20 mins | |
| 11:25 | General 4: **Adaptation** 7 talks | General 2: **Biodiversity** 7 talks |
| 13:10 | Lunch * 1.05 hours | |
| 13:30 | Lunch Time Social Event | |
| 14:15 | **Keynote: Stephane Hourdez** | |
| 14:50 | General 4: **Adaptation** 3 talks & Special 6a: **Biomimicry** 5 talks | General 2: **Biodiversity** 8 talks |
| 16:50 | Coffee Break * 20 mins | |
| 17:10 | General 6: **Biomimicry** Round table | Special 1d: **Stewardship** Round table & 4 talks |
| 19:30 | **Online only:** Annual General Meeting (AGM) DSBS | |

| **Friday 17/09** | | |
|---|---|---|
| 08:30 | **Keynote: Kim Juniper** | |
| 09:05 | Special 5b: **Artificial Intelligence** 6 talks & **General 5: Access** 2 talks | General 3: **Life history & connectivity** 8 talks |
| 11:05 | Coffee Break * 20 mins | |
| 11:25 | General 5: **Access** 2 talks & **Special 5a: Ecosystem dynamics** 5 talks | General 3: **Life history & connectivity** 7 talks |
| 13:10 | Lunch * 1.05 hours | |
| 13:30 | Lunch Time Social Event | |
| 14:15 | **Tribute to Anne Rognant** | |
| 14:30 | Special: **Arts & science** 6 talks clotured by a music event (no coffee break) | General 2: **Biodiversity** 6 talks |
| 16:00 | Coffee Break * 20 mins | |
| 16:20 | Special: **Open session** 4 talks | Special 1a: **UN BBNJ** Round table |
| 17:30 | Closing: Prize awards, results of the voting for the next Symposium venue | |
